# Binaural Beats through the auditory pathway: from brainstem to connectivity patterns

**DOI:** 10.1101/623231

**Authors:** Hector D Orozco Perez, Guillaume Dumas, Alexandre Lehmann

## Abstract

Binaural beating is a perceptual auditory illusion occurring when presenting two neighboring frequencies to each ear separately. Binaural beats have been attributed to several controversial claims regarding their ability to modulate brain activity and mood, in both the scientific literature and the marketing realm. Here, we sought to address those questions in a robust fashion using a single-blind, sham-controlled protocol. To do so, we characterized responses to theta and gamma binaural beats and “sham” stimulation (monaural beats) across four distinct levels: subcortical and cortical entrainment, scalp-level Functional Connectivity and self-reports. Both stimuli elicited standard subcortical responses at the pure tone frequencies of the stimulus (i.e., Frequency Following Response), and entrained the cortex at the beat frequency (i.e., Auditory Steady State Response). Furthermore, Functional Connectivity patterns were modulated differentially by both kinds of stimuli, with binaural beats being the only one eliciting cross-frequency activity. Despite this, we did not find any mood modulation related to our experimental manipulation. Our results provide evidence that binaural beats elicit cross frequency connectivity patterns, but weakly entrain the cortex when compared to a sham stimulus. Whether these patterns have an impact in cognitive performance or other mood measurements remains to be seen.

**Significance Statement:** Binaural beats have been a source of speculation and debate in the scientific community. Our study addresses pseudo-scientific marketing claims and approaches them using proper experimental control and state-of-the-art signal processing techniques. Here we show that binaural beats can both entrain the cortex and elicit specific connectivity patterns. Regardless of this, our sham condition was able to entrain the cortex more strongly, and both binaural beats and the sham condition failed to regulate mood. All in all, though binaural beats weakly entrain cortical activity and elicit complex patterns of connectivity, the functional significance (if any) of these patterns remains an open question.

## Introduction

Humans use music and rhythm as mood enhancers. Be it in social gatherings or late study nights, we use audio stimuli to set the “right mood” and improve our cognitive performance. Binaural beats, an auditory illusion that occurs when presenting two similar tones to each ear separately, have been advertised to induce mood alterations dependent on the beat frequency. Claims range from entraining the whole brain (Atwater, 2004; Rhodes, 1993), to the alteration of states of consciousness (I-Doser, accessed May 2018; Atwater, 1997). The possibility of binaural beats entraining and modulating cortical activity without previous training makes them an interesting candidate for mood regulation applications and off-the-shelf cognitive enhancement.

Presenting two carrier tones with a slight frequency mismatch to each ear separately creates a perception of a third tone, a binaural beat, that oscillates at the absolute difference between the tones (Oster, 1973; Moore, 2012). These beats are thought to originate subcortically in the medial nucleus of the superior olivary complex, the first nucleus in the auditory pathway to receive bilateral input (Wernick & Starr, 1968; Kuwada & Wickesberg, 1979). Binaural beats can entrain cortical activity at both the specific frequency of the beat (Draganova et al., 2007; Schwarz & Taylor, 2005; Pratt et al., 2010) and cross-frequency modulations, such as theta beats driving interhemispheric alpha synchronization (Solcà et al., 2016). They also seem to modulate mood (Wahbeh et al., 2007; Le Scouarnec et al., 2001; Padmanabhan et al., 2005), pain perception (Zampi, 2015) and cognitive performance in memory tasks (Lane et al., 1998; Kennerly, 1994; Garcia-Argibay, 2017). These results, however, seem to be inconsistent and dependent on several mediating factors, such as frequency of stimulation, data analysis discrepancies, exposure time and stimuli masking (Garcia-Argibay et al., 2018). Furthermore, no study to date has fully characterized binaural beats throughout the auditory pathway (from subcortical responses to Functional Connectivity) and compared their effect to that of a rhythmic sham control. Indeed, isochronic tones readily entrain the cortex to specific frequencies (Nozaradan et al., 2016), and repetitive and rhythmic stimuli (such as mantras or tones) are widely used in contemplative and religious practices with positive physiological impact (Bernardi et al., 2001; Bernardi et al., 2017). Whether binaural beats are actually a special kind of stimulus, and if their effects are due to their rhythmicity, or a placebo effect, remain open questions. To address these, we recorded electroencephalography (EEG) during a single-blind, sham-controlled task in which participants passively listened to either binaural beats or monaural beats (the rhythmic sham condition, created by digitally summing each tone before presentation).

Our objective was threefold: to fully characterize brain and mood responses to binaural beats, to compare those to monaural beats (sham condition) and to compare the effect of two different beat frequencies. We used theta (7 Hz) and gamma (40 Hz) beats because theta beats have been associated with reduced anxiety levels (McConnell et al. 2014; Isik et al., 2017), and gamma beats have been associated with attention modulation (Colzato et al., 2017). Furthermore, these two frequencies have been associated with large-scale integration models of the brain (Varela et al., 2001; Canolty and Knight, 2010). We compared responses between binaural and monaural beats at four levels: subcortical entrainment to the carrier tones in the form of a Frequency Following Response (FFR; Skoe & Kraus, 2010), cortical entrainment to the beat in the form of an Auditory Steady State Response (ASSR; Picton et al., 2003), changes in Functional Connectivity using phase-based statistics (Nolte et al. 2004; Lachaux et al., 2000) and mood changes as self-reported using a visual analogue scale (Rainville et al., 2002). We hypothesized that binaural and monaural beats would both elicit cortical and subcortical responses to the beat (ASSR) and pure tone frequencies (FFR) respectively. However, we expected that binaural beats would elicit Functional Connectivity changes and modulate mood, with no such changes during the sham condition. Furthermore, we hypothesized that theta beats would facilitate a peaceful and relaxed state, while gamma beats would elicit a more alert state. As Solcà et al. (2016) imply, using these sequential levels allowed us to assess the robustness of our observations. By presenting converging evidence from different approaches (self-reports, EEG cortical and subcortical entrainment and connectivity patterns), we aimed to elucidate whether binaural beats are actually a special kind of stimulus, or reported effects are in part due to their rhythmicity.

## Materials & Methods

To understand the functional meaning of the entrainment and connectivity patterns associated with binaural beats, we investigated the differences between monaural and binaural beats by comparing subcortical, cortical and subjective responses elicited through a blind, passive listening task with a 2×2 factorial design (two within factors: beat type and beat frequency).

### Participants

Sixteen participants (nine female, seven male; mean age 27.4±5.5) volunteered for the experiment and provided written informed consent. Exclusion criteria included neurological damage or abnormalities (e.g. demyelination), and major hearing loss (0-20 HL dB) as self-reported by the participants. The experimental procedures conformed to the World Medical Association’s Declaration of Helsinki and were approved by the Research Ethics Committee of the Faculty for Arts and Sciences of the University of Montreal.

### Stimuli

Binaural beat stimuli consisted of two pure sine tones with equal starting phase and a slight frequency mismatch presented separately to each ear (**Figure 1, Columns 1 and 2**). These two pure tones were superimposed digitally to create the sham condition, which was presented to both ears. We refer to beat frequency as the difference between these sine waves; either 40 Hz for gamma conditions (380 Hz & 420 Hz) or 7 Hz for theta conditions (396.5 Hz & 403.5 Hz). We chose carrier frequencies around 400 Hz for two reasons: best perception of binaural beats occurs at carrier tones between 400 and 500 Hz (Licklider et al., 1950; Perrot & Nelson, 1969), and this frequency range minimizes cortical contributions to the brainstem responses (Coffey et al., 2016). Both kinds of stimuli (binaural and sham) were root mean squared normalized.

**Figure 1.**
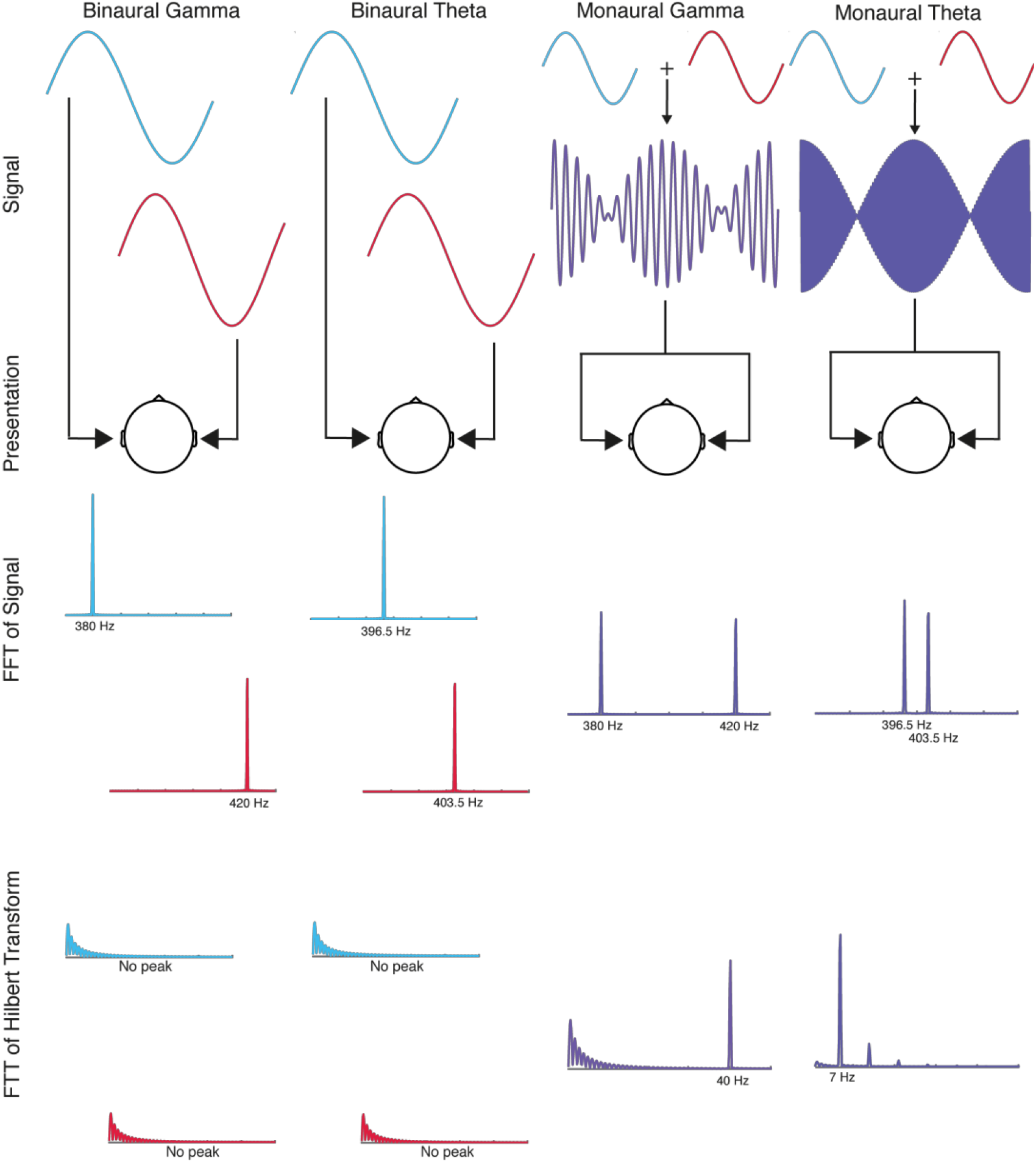
Beats: Signal, Presentation method, Fast Fourier Transform (FFT) and FFT of Hilbert Figure Transform. Each column represents one experimental condition. (**Signal and Presentation rows**) Binaural beats are created by dichotically presenting two pure tones with a slight frequency mismatch (Red color = right ear). Monaural beats are created by digitally summing these tones and presenting the resulting signal diotically. (**FFT of Signal**) Stimuli were analyzed using a Fourier transform to obtain their frequency composition. (**FFT of Hilbert Transform**) The FFT of the Hilbert transform (i.e. the analytic signal) was computed to tap into the spectral information of the envelope of the signal (the beat frequency). The frequency of the envelope of the summed tones encodes beat frequency (e.g. 403.5 - 396.5 = 7 Hz for Theta). This information, however, is only encoded in monaural beats because they are digitally summed.

#### Frequency choice: Theta

Auditory stimulation at the theta frequency band (4 - 7 Hz) has been associated with positive emotional experiences and introspection (Aftanas & Golocheikine, 2001), reduced perceived pain in patients with chronic pain (Zampi, 2015), states of meditation and decreased alertness (Jirakittayakorn & Wongsawat, 2017) and enhancement of immediate verbal memory (Ortiz et al., 2008). Furthermore, theta cortical activity is related to concentration, focused attention and a general meditative state (Takahashi et al., 2005; Lagopoulos et al., 2009). We chose theta beat frequency to explore the possibility of eliciting a mindful and relaxed state in the participants.

#### Frequency choice: Gamma

Auditory gamma stimulation (32 - 48 Hz) has been associated with binaural sound integration (Ross et al., 2014), divergent thinking (Reedijk et al., 2013), and attention control (Reedijk et al., 2015). Furthermore, auditory cortices readily are entrained to it (Schwarz & Taylor, 2005; Ross et al., 2014), and it seems to be a “natural frequency” of these areas, even during resting state (Hillebrand et al., 2012). We chose gamma beat frequency to explore the possibility of eliciting a heightened attention cognitive state in the participants.

### Procedure

Participants started by filling out a general information and music abilities questionnaire. We then fitted a headcap on participants’ heads and placed EEG electrodes in it using conductive gel. The experiment took place in a sound-attenuated, electromagnetically shielded room at the International Laboratory for Brain, Music and Sound Research (BRAMS; Montreal, Canada). Participants were asked to relax their upper body, close their eyes, avoid body movements and to pay attention to the beat throughout the experiment (Schwarz et al., 2005). We recorded data from five experimental blocks: an eight minute baseline (no stimulus presentation; eyes-closed) followed by the four pseudo-randomized experimental conditions (binaural gamma, monaural gamma, binaural theta, monaural theta), each lasting for eight minutes. After each recording block, participants were asked to rate their experience using two visual analogue scales. They were also given the opportunity to take a break in the middle of the experiment. Auditory stimuli (both binaural and monaural beats) were generated live (i.e. during the recording block) to ensure sub-millisecond phase accuracy using a signal processing system (RX6, Tucker-Davis Technologies, www.tdt.com) controlled with MATLAB software (The Mathworks, www.mathworks.com) and delivered via insert earphones (ER3, Etymotic Research, www.etymotic.com). Auditory stimuli were processed at 48 kHz and were each presented continuously for eight minutes at 70 dB SPL. For the purpose of further analysis and the epoching of continuous data, triggers were sent every eight seconds via parallel ports using the signal processing system (RX6, Tucker-Davis Technologies, www.tdt.com) and recorded along with the EEG data.

#### EEG data acquisition

EEG was recorded using 64 active sintered Ag-AgCl electrodes placed on the scalp according to the International 10/10 system (ActiveTwo, Biosemi, The Netherlands). The active electrodes contain the first amplifier stage within the electrode cover and provide impedance transformation on the electrode to prevent interference currents from generating significant impedance-dependent nuisance voltages. We, therefore, did not control electrode impedances but rather kept direct-current offset close to zero during electrode placement. Vertical and horizontal eye movements were monitored using three additional electrodes placed on the outer canthus of each eye and on the inferior area of the left orbit. Reference-free electrode signals were amplified, sampled at 2048 Hz (ActiveTwo amplifier, BioSemi, The Netherlands), and stored using BioSemi ActiView Software for offline analysis. Given that auditory stimuli were created online during the experiment, they were recorded using Biosemi’s Analog Input Box (Biosemi, The Netherlands), which was daisy chained by optical fibers to the EEG Analog-to-Digital Converter box and stored alongside the EEG data for future analysis.

#### Visual Analogue Scales

Participants were given pen and paper analogue scales after each recording block so they could rate their experience after the passive listening task. Two analogue scales were used to determine variations in subjective experience (Rainville et al., 2002). The scales used were:

- Mental relaxation, corresponding to the activity or calmness of the subject’s mind. This dimension spans from a state where the mind is calm, peaceful and in perfect relaxation to a state where the mind is extremely agitated or active.
- Absorption depth corresponds to how the subject feels and how absorbed they felt during the experiment. The scale runs from nonexistent depth to a profound, intense and complete experience.

### Data analysis and signal processing

#### Visual Analogue Scales

Data from pen and paper scales was measured manually and stored digitally in CSV files for further statistical analysis using R (v3.2.3, R Development Core team, 2008), setting the significance level at 0.05. We analyzed the data using a one-way repeated measures ANOVA for each scale (Mental Relaxation and Absorption Depth) with “Condition” as a 5-level factor (Baseline, Monaural Gamma, Binaural Theta, Binaural Gamma, Monaural Theta). We used *post-hoc* paired t-tests to further disentangle patterns in the data. We kept the Family-wise error rate (FWER) at p = 0.05 by using Holm’s sequential Bonferroni procedure.

#### EEG

The data was processed using the EEGLAB toolbox (Delorme A. & Makeig S., 2004) and in-house developed scripts in MATLAB. Two different analyses were conducted on the raw EEG data: subcortical (Frequency Following Response) and cortical (Auditory Steady State Response and Functional Connectivity). The pre-processing procedures for either subcortical or cortical analysis differed in: filtering process, ICA decomposition and re-referencing. For subcortical analysis, data was high-pass filtered at 100 Hz and re-referenced to linked mastoids. Data used for cortical analysis was bandpass filtered between 1-100 Hz, decomposed using ICA for artifact correction purposes (Jung et al., 1998) and re-referenced to linked mastoids as a first step and then to common average reference as a final step.

#### Frequency Following Response (FFR)

Data was re-referenced to linked mastoids and high-pass filtered at 100 Hz using a zero-phase Butterworth filter Order 4. Data was visually inspected for noisy electrodes, which were then removed and interpolated using spherical interpolation. Finally, data was epoched into 60 events (from −1 to 7 s with respect to trigger onset) and exported for further analysis. Epochs from each participant were averaged and transformed into the frequency domain using an FFT. From these, power was calculated as the square of the magnitude normalized using a factor of *2/N, N* being the length of the epoch. Frequencies of interest were extracted as the mean of 1 Hz bins around the carrier frequencies (pure tones: 380, 397.5, 403.5, and 420 Hz) for the baseline and each experimental condition. A baseline normalization (decibel change from baseline) was performed to disentangle background dynamics from actual stimulation-related oscillations (Cohen, 2014). The equation used was as follows:

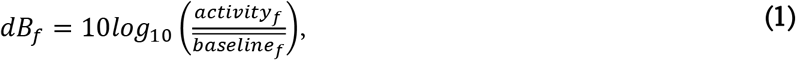

where *activity_f_* is a specific frequency power in a given experimental condition and 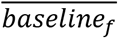 is the average activity across the whole baseline at a given frequency (Cohen, 2014). The unit of this data is decibel change from baseline.

After baseline normalization, all the scalp channels were averaged together to output one normalized power score per experimental condition per participant. Frequency relevant scores were averaged together in each experimental condition. For example, power scores for 396.5 and 403.5 Hz were averaged together for theta frequency relevant scores. This was done in order to keep the hypothesis testing at a minimum and avoiding inflating the family-wise error rate (FWER). These averaged scores were then exported to R (v3.2.3, R Development Core team, 2008) for hypothesis testing.

Two sets of data were analyzed (power at relevant gamma and relevant theta frequencies) with 64 scores each (4 conditions x 16 participants) for statistical significance. A factorial (2×2) repeated measures ANOVA was computed per relevant pure tone data set (Theta and Gamma) using beat type (binaural, monaural) and frequency (gamma, theta) as factors. Partial omega-squared is reported as a measurement of effect size. To further disentangle patterns in the data, *post-hoc* paired t-tests using Holm’s sequential Bonferroni correction were used (as to keep the FWER at 0.05).

#### Auditory Steady State Responses (ASSR)

Data was imported and re-referenced to linked mastoids. Data was then resampled at 512 Hz, trimmed around the time-window of interest (8m ± 3s) and filtered twice: using a 2nd order Butterworth band-pass filter (zero-phase) between 1-100 Hz, and an FIR notch filter at 60 Hz (minimizing line noise). Data was visually inspected for noisy electrodes, which were then removed.

For each participant, ICA decomposition was applied to the full recording of all conditions. Prior to this, these aggregated files were first filtered between 1 and 80 Hz and decimated to 256 Hz. Data was then decomposed using the *runica()* function from EEGlab (which uses the Bell & Sejnowski’s (1995) ICA algorithm and Lee, Girolami & Sejnowski’s extended-ICA algorithm (2000)). After visual inspection of individual components, weight matrices were obtained from this decomposition and applied to the original five files for artifact correction purposes (remove components deemed as non-cortical activity, Jung et al., 1998). Missing electrodes were interpolated after ICA artifact correction. EEG was re-referenced to common average (CAR) and epoched from - 1s to 8s relative to trigger onset. Finally, data was baseline corrected (using whole epoch as the baseline) and stored for further analysis.

Each participant’s epochs were averaged and transformed into the frequency domain using an FFT. From these, power was calculated as the square of the magnitude normalized using a factor equal to 2/*N*, where *N* is the number of samples in each sequence. Frequencies of interest were extracted as the mean of 1 Hz bins around beat frequencies (7 and 40 Hz) for both the baseline and each experimental condition. As with the FFR preprocessing procedure, a baseline normalization (decibel change from baseline) was done using **Equation 1**.

After baseline normalization, all channels were averaged together to output one normalized power score per experimental condition per participant. These scores were then exported to R (v3.2.3, R Development Core team, 2008) for hypothesis testing.

Statistical analysis were very similar to those done for the FFR analysis. Two statistical analysis were performed: one on normalized power scores at gamma beat frequency at each experimental condition, and one on normalized power scores at theta beat frequency at each experimental condition (each had 64 scores total, 4 conditions x 16 participants). Hypothesis testing was performed using a factorial (2×2) repeated measures ANOVA with beat type (binaural, monaural) and frequency (gamma, theta) as factors. Partial omega-squared is reported as a measurement of effect size. Finally, *post-hoc* paired t-tests using Holm’s sequential Bonferroni procedure to keep the Family-wise error rate (FWER) at 0.05 were used to further disentangled patterns in the data.

#### Functional Connectivity

Two complementary measurements of Functional Connectivity were used as indices of long-range synchronizations: Phase locking value (PLV; Lachaux et al., 2000)) and Imaginary Coherence (iCOH; Nolte et al., 2004). On top of that, per electrode, the amplitude of the Hilbert Transform and the Power of the Fourier transform were computed as local indices of synchronization. Analyses were done over all traditional frequency bands (Delta: 1 - 4 Hz; Theta: 5 - 8 Hz, Alpha: 9 - 12 Hz, Beta: 13 - 30 Hz, Gamma: 32 - 48 Hz) and the specific beat frequencies (1Hz bins around 7Hz and 40 Hz). ICA corrected data (i.e. the same files used for ASSR) was imported to MATLAB to compute these metrics.

#### Hilbert Transform and PLV

The phase locking value (PLV) looks at how stable phase differences are between signals (in this case, electrodes). In this particular implementation, it determines, on average, how stable phase differences between electrodes are within trials. PLV is only sensitive to phase differences between signals (not their amplitude) at the cost of not being able to distinguish spurious correlation due to volume conduction at the scalp level from actual connectivity between two cortical regions (Lachaux et al., 1999).

To calculate it, the signal of interest was extracted by band-passing the ICA corrected data using a finite impulse response filter (FIR) around both the traditional frequency bands of interest and the specific beat frequencies: Delta (1 - 4 Hz), Theta (5 - 8 Hz), Alpha (9 - 12 Hz), Beta (13 - 30 Hz), Gamma (32 - 48 Hz), Theta Beat (6 - 8 Hz) and Gamma Beat (39 - 41 Hz). Phase and amplitude of the analytical signal (Hilbert transform) were then extracted for each EEG channel. For each pair of electrodes, the PLV was computed as a long-distance synchronization index on eight-second non-overlapping sliding windows as

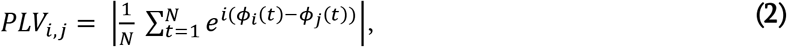

where N is the number of samples considered in each eight second window, *ϕ* is the phase and || the complex modulus. Thus, PLV measure equates 1 if the two signals are perfectly phase locked across the whole observed time window, and equates 0 if they are totally unsynchronized. For each electrode, the amplitude of the analytic signal (Hilbert Transform) was stored as a local synchronization index. Nonparametric permutation testing was used to gauge the statistical significance of the effects of binaural and monaural beats on Functional Connectivity.

#### Fourier Transform and iCOH

Coherency (Magnitude-Squared Coherence) between two EEG channels can be defined as the measure of a linear relationship (i.e. correlation) between two signals (in this case, electrodes) at specific frequencies. It is calculated as the cross-spectral density between channels *i* and *j*, normalized by the square root of the multiplication of each of their own auto-spectrums. By projecting the results into the imaginary axis, we rid the signal of both immediate (a phase difference of 0) and anti-phase (phase difference of π) connectivity patterns. The imaginary part of coherence is insensitive to spurious correlations due to volume conduction at the expense of being sensitive to signals’ amplitude (as well as phase) and being unable to disentangle spurious from real immediate connectivity patterns (both in phase and anti-phase; Nolte et al. 2004).

Imaginary coherence measures were extracted on eight-second non-overlapping sliding windows (similar to the PLV procedure):

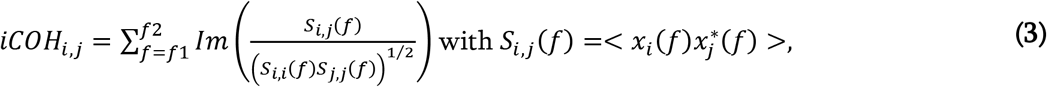

where *x_i_*(*f*) and *x_j_*(*f*) are the complex Fourier transforms of channels *i* and *j* respectively, * stands for complex conjugation, <> for the expectation value, *f*_1_ and *f*_2_ are the boundary of the considered frequency band, *S_i,j_*(*f*) is the cross-spectral density between channels *i* and *j*, and *Im*() is the imaginary part of a complex number. As a long-distance synchronization index, iCOH values were averaged for each pair of electrodes across frequency bins using a tolerance of 1 Hz (e.g., 7 ± 1 Hz). Each electrode’s autospectrum was stored as a local synchronization index. As with PLV and Hilbert Transform, nonparametric permutation testing was used to gauge the statistical significance of the effects of binaural and monaural beats on Functional Connectivity.

#### Cross-frequency interactions

In this context, we consider cross-frequency interactions as activity elicited by either experimental condition (binaural or sham) that is outside of the frequency range of the beat (either 7 Hz for theta, or 40 Hz for gamma). For example, activity in the alpha frequency band elicited by theta experimental conditions is considered as a cross-frequency interaction (Solcà et al., 2016).

#### Neurophenomenological analysis

To explore the relationship between mood (as self-reported by the visual analogue scales) and neural patterns of Functional Connectivity, each participant’s two highest rated experimental conditions (binaural gamma, monaural theta…) were contrasted with the two lowest rated ones for each visual analogue scale. These contrasts were then averaged across participants for each scale (Mental Relaxation and Absorption Depth). Again, non-parametric permutation testing was used to gauge the statistical significance of the relations between subjective experience and Functional Connectivity.

#### Functional Connectivity: Nonparametric statistics

Given the exploratory nature of our study, we decided to use nonparametric permutation testing to maintain the family-wise error rate (FWER) at 5%, as it offers a straightforward solution to the multiple-comparisons problem (see Maris & Oostenveld, 2007; and Groppe et al., 2011). The critical *t* value was determined for all Functional Connectivity analysis (PLV, iCOH, autospectrum, Hilbert transform, and neurophenomenology) as follows: (1) the experimental conditions were contrasted with each other (binaural vs. sham) and each experimental condition was contrasted with baseline, (2) a *t*-test was performed at each spatial-spectral point (i.e. electrode at a given frequency), (3) the statistics were normalized using z-scores, (4) the cluster statistic was considered to be the sum of all *t*-values of the cluster members exceeding 3 in absolute value, (5) 1,000 permutations of the data were then performed to obtain a distribution of cluster statistics under the null hypothesis and determine the critical values. All randomizations were done for a rejection of the null hypothesis and a control of false alarm rate at p < 0.05. We decided to choose this method to correct for multiple comparisons because we are mainly interested in broadly distributed effects (Groppe et al., 2011). To make our inferences more conservative, only contrasts that exhibit at least three significant spatio-spectral points are shown here (i.e. electrodes at a given frequency; contrasts with no significant patterns or with less than three points can be seen in **Supplemental Figures S2 - S7**).

## Results

### Visual Analogue Scales

There was no effect from auditory stimulation on subjective experiential ratings. There were no differences between the five levels of the factor Condition (baseline, binaural gamma, binaural theta, monaural gamma, monaural theta) neither in Mental Relaxation (F(4, 60)=1.698, p = .162) nor Absorption Depth (Mauchly’s test revealed a violation of the assumption of sphericity; *p* values corrected using Greenhouse-Geisser estimates: F(4, 60)=1.313, p_corr_ = 0.284, *ε*= 0.537). These suggest that subjective experience related to each experimental condition was not different from baseline nor from each other (**Supplemental Figure S1**).

### Frequency Following Response (FFR)

#### Theta Pure Tones (396.5 & 403.5 Hz)

Both theta binaural and monaural beats elicited an FFR at theta carrier frequencies with no difference between them (average of 396.5 Hz and 403.5 Hz; **Figure 2a**). There was a main effect of beat frequency (F(1, 15) = 34.573, p = 0.00003, 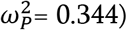), with no effect of beat type (F(1, 15) = 0.0040, p = 0.95) nor an interaction between the factors (F(1, 15) = 1.1687, p = 0.2976).

**Figure 2.**
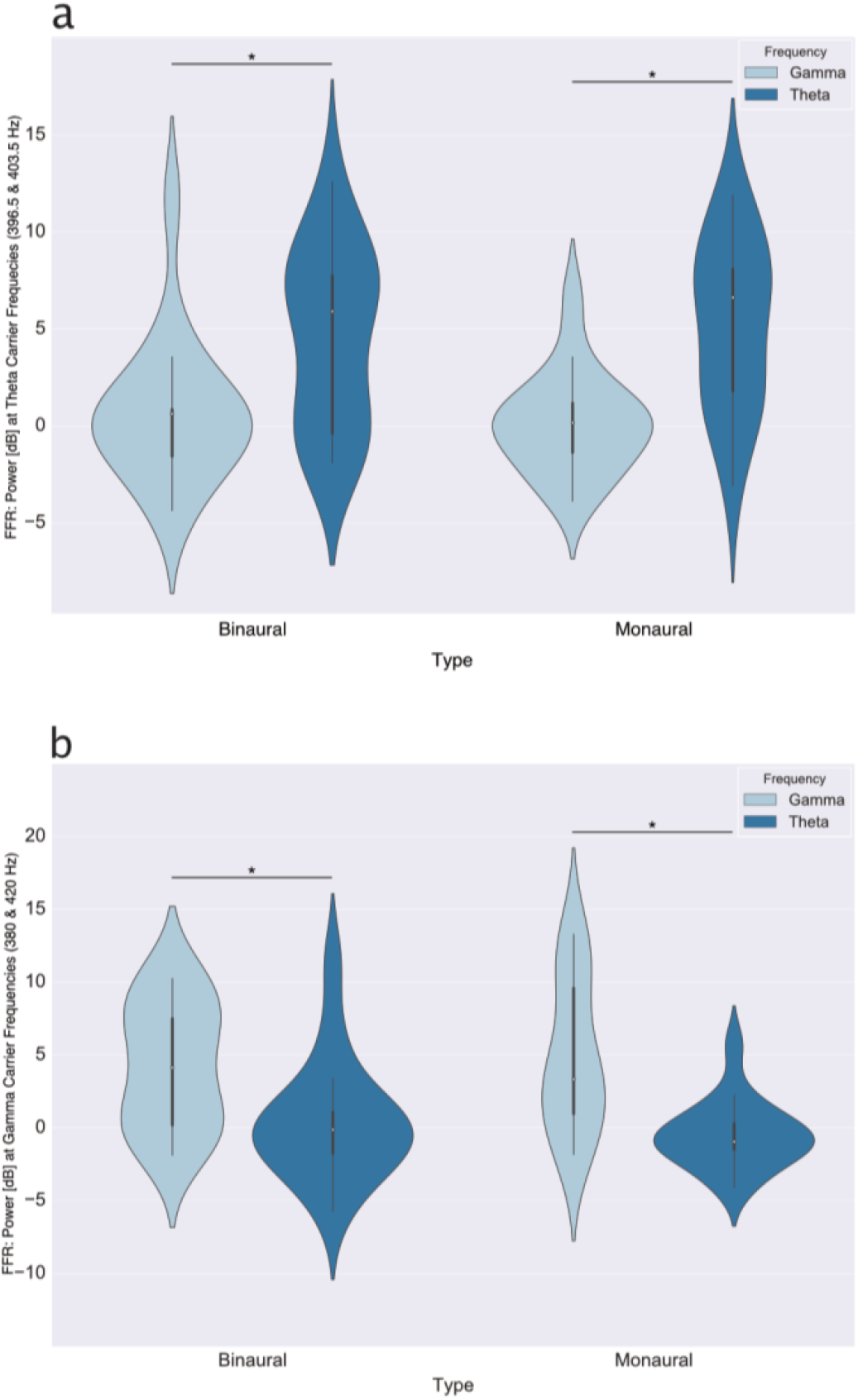
Frequency Following Response (FFR) to carrier pure tones. Plotted here: violin plots with median (white dots), quartile (thick black line) and whisker (thin black line) values. Please note the scale is decibel change from baseline, a logarithmic scale where each 3 dBs represent a difference of a factor of 2. Each violin plot represents all participants’ baseline-normalized (dB) averages of the power around a 1 Hz bin (e.g. 396.5 ± 0.5 Hz) at beat carrier frequencies (e.g. 396.5 and 403.5 Hz were averaged together for theta conditions). This power was obtained from the average activity at all channels of each participant. Asterisks above lines linking conditions denote a significant difference between them (p < 0.05). **(a)** Frequency Following Response elicited at theta-carrier frequencies (average of 396.5 and 403.5 Hz). **(b)** Frequency Following Response elicited at gamma-carrier frequencies (average of 380 and 420 Hz).

#### Gamma Pure Tones (380 & 420 Hz)

Gamma binaural and monaural beats elicited an FFR at gamma carrier frequencies (average of 380 Hz and 420 Hz; **Figure 2b**), with no difference between them. There was a main effect of frequency (F(1, 15) = 26.648, p = 0.00012, 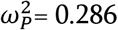) with no effect of beat type (F(1, 15) = 0.0565, p = 0.8152) nor an interaction between the factors (F(1, 15) = 1.2476, p = 0.2815).

### Auditory Steady State Response (ASSR)

#### Theta ASSR (7 Hz)

Theta binaural and monaural conditions elicited an ASSR at beat frequency, with monaural beats peaking higher than binaural beats (**Figure 3a**). There was both a main effect of beat type (F(1, 15) = 7.6686, p = 0.01432, 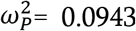) and beat frequency (F(1, 15) = 19.2633, p = 0.00053, 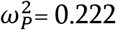), with no interaction (F(1, 15) = 3.928, p = 0.0661).

**Figure 3.**
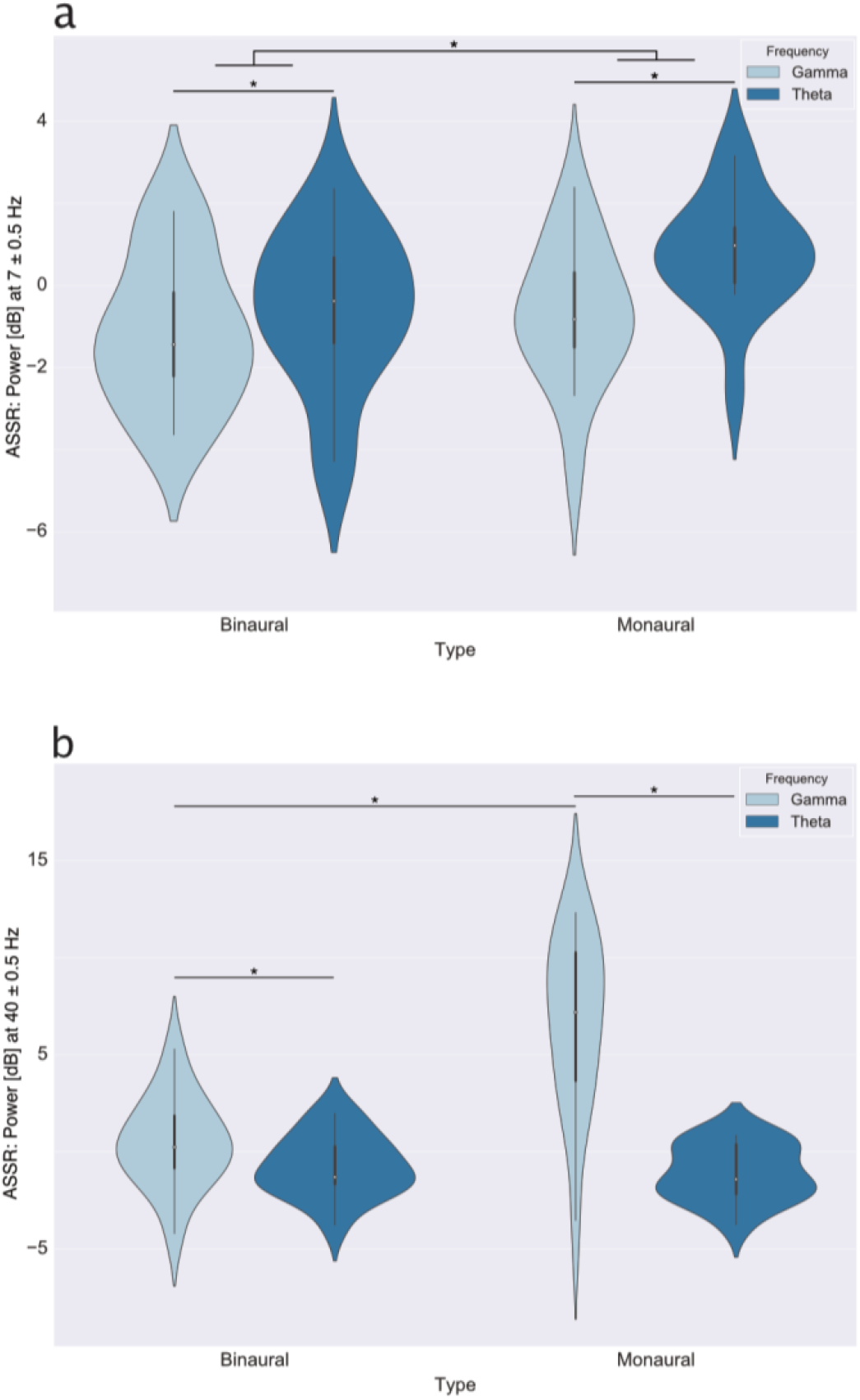
Auditory Steady State Responses (ASSR) to beats. Plotted here: violin plots with median (white dots), quartile (thick black line) and whisker (thin black line) values. Please note the scale is decibel change from baseline, a logarithmic scale where each 3 dBs represent a difference of a factor of 2. Each violin plot represents all participants’ baseline-normalized (dB) averages of the power around a 1 Hz bin (e.g. 7 ± 0.5 Hz) at beat frequencies (either 7 or 40 Hz) obtained from the average activity at all channels for each participant. Asterisks above lines linking conditions denote a significant difference between them (p < 0.05). Please note that there was an outlier in these graphs that was taken out for visualization purposes (a participant with data points at around, 30 dB). **(a)** Cortical activity elicited at 7 Hz. **(b)** Cortical activity elicited at 40 Hz.

#### Gamma ASSR (40 Hz)

Gamma beats (both binaural and monaural) elicited an ASSR at 40 Hz, with binaural gamma eliciting the highest power (**Figure 3b**). There were main effects of both beat type (F(1, 15) = 34.5383, p = 0.00003, 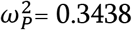) and beat frequency (F(1, 15) = 51.9329, p = 0.000003, 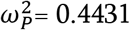), as well as an interaction between the two factors (F(1, 15) = 44.2841, p = 0.000007, 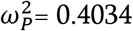). To further disentangle these differences, three post-hoc pairwise comparisons were done using Holm’s sequential Bonferroni correction test. The first two comparisons confirmed that monaural gamma peaked the highest at 40 Hz when compared to both binaural gamma (Mean difference = 6.2702, p_corr_ = 0.000005) and monaural theta (Mean Difference = 7.6589, p_corr_ = 0.000004). Binaural gamma condition elicited a stronger ASSR than binaural theta (Mean Difference = 1.1888, p_corr_ = 0.02637).

### Functional Connectivity

#### Phase-locking value (PLV) and Hilbert Transform Amplitude

Both binaural and sham conditions elicited within and cross-frequency patterns at long and short ranges. These were dependent on both beat type and frequency. In terms of local synchronization (Hilbert Transform Amplitude), Monaural Gamma stimulation drove a positive frontoparietal cluster at 40 Hz (gamma beat) when contrasted with baseline (**Figure 4a**, CS = 20.52, p = 0.026). In terms of long-distance synchronization (PLV), we found a positive left-occipital to frontoparietal cluster of activity at 40 Hz (Figure 4b, CS = 49.83, p = 0.036) when contrasting binaural theta with monaural theta experimental conditions. When contrasting binaural gamma with monaural gamma, we found four clusters of activity: a positive cluster extending around the scalp at alpha frequency band (**Figure 4c top left**, CS = 1043.45, p = 0.004), a negative central-occipital cluster at gamma frequency band (**Figure 4c top right**, CS = −219.57, p = 0.043); a negative frontal cluster at gamma frequency band (**Figure 4c center**, CS = −240.17, p = 0.028); and a negative scalp-wise cluster at 40 Hz (**Figure 4c bottom left**, CS = −2695.07, p = 0.003). Consistent with the last cluster, the monaural condition drove a positive scalp-wise cluster when contrasted with baseline (**Figure 4c bottom right**, CS = 2493.34, p=0.006). All Hilbert Amplitude and PLV contrasts (both significant and non-significant) can be found in the supplemental material (**Figure S2** for Theta conditions and **Figure S3** for Gamma conditions).

**Figure 4.**
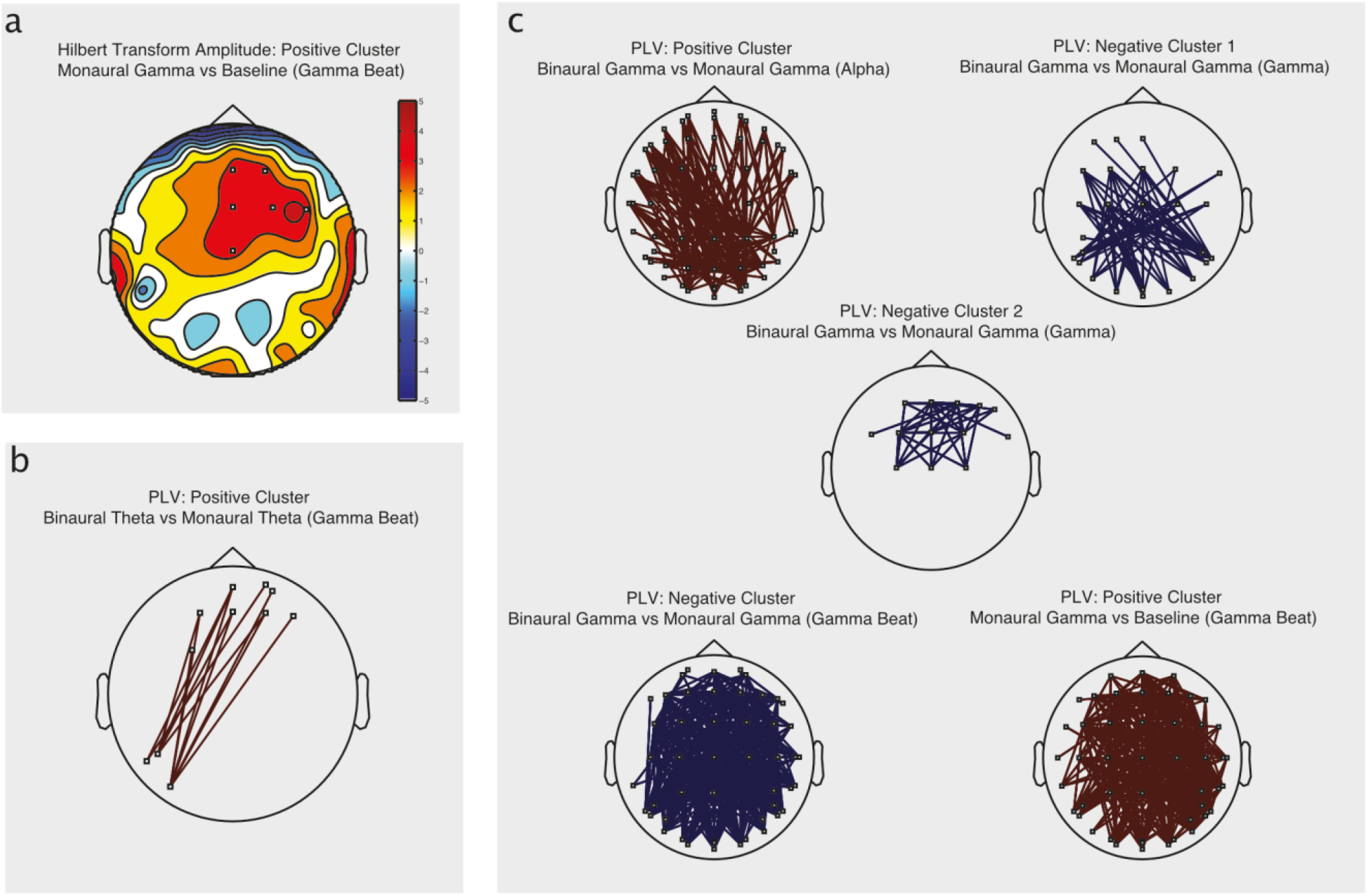
Contrast topographies for Phase-locking value (PLV) and Hilbert Transform amplitude. Topographies were averaged across participants and compared to either baseline or between beat type (binaural vs monaural). Both statistics (Hilbert Transform Amplitude and PLV) were normalized using z scores. We only show contrasts that exhibit at least three significant electrodes here (depicted as small white squares). All contrasts can be found in the supplemental material (refer to **Figure S2** (Theta conditions) and **Figure S3** (Gamma conditions)). Frequency band limits are as follows: Delta (1 - 4 Hz), Theta (5 - 8 Hz), Alpha (9 - 12 Hz), Beta (13 - 30 Hz), Gamma (32 - 48 Hz), Theta Beat (6 - 8 Hz) and Gamma Beat (39 - 41 Hz). **(a) Hilbert transform amplitude used as a local synchronization index.** The color bar indicates T-values from student’s test. **(b) Phase locking value used as an index of long-distance synchronization between electrodes during theta conditions.** Red lines indicate a significant positive phase locking value (PLV) between two electrodes. **(c) Phase locking value used as an index of longdistance synchronization between electrodes during gamma conditions.** Red lines indicate a significant positive phase locking value (PLV) between two electrodes while blue lines indicate a negative one.

#### Imaginary Coherence (iCOH) and Fourier Transform Power

As indexed by iCOH and Fourier Transform, we only found short distance synchronization elicited by binaural theta conditions. In terms of local synchronization (Fourier Power), we found two clusters of activity when contrasting binaural theta condition with baseline: a negative central-parietal cluster of activity at theta frequency band (**Figure 5, top**; CS = −11.45, p = 0.044) and a positive left central-temporal cluster at 40 Hz (**Figure 5 bottom**; CS = 36.10, p = 0.013). None of the other contrasts reached our criteria for significance (p < 0.05 and a cluster of at least three sensors). All Fourier Power and iCOH contrasts (both significant and non-significant) can be found in the supplemental material (**Figure S4** for Theta conditions and **Figure S5** for Gamma conditions).

**Figure 5.**
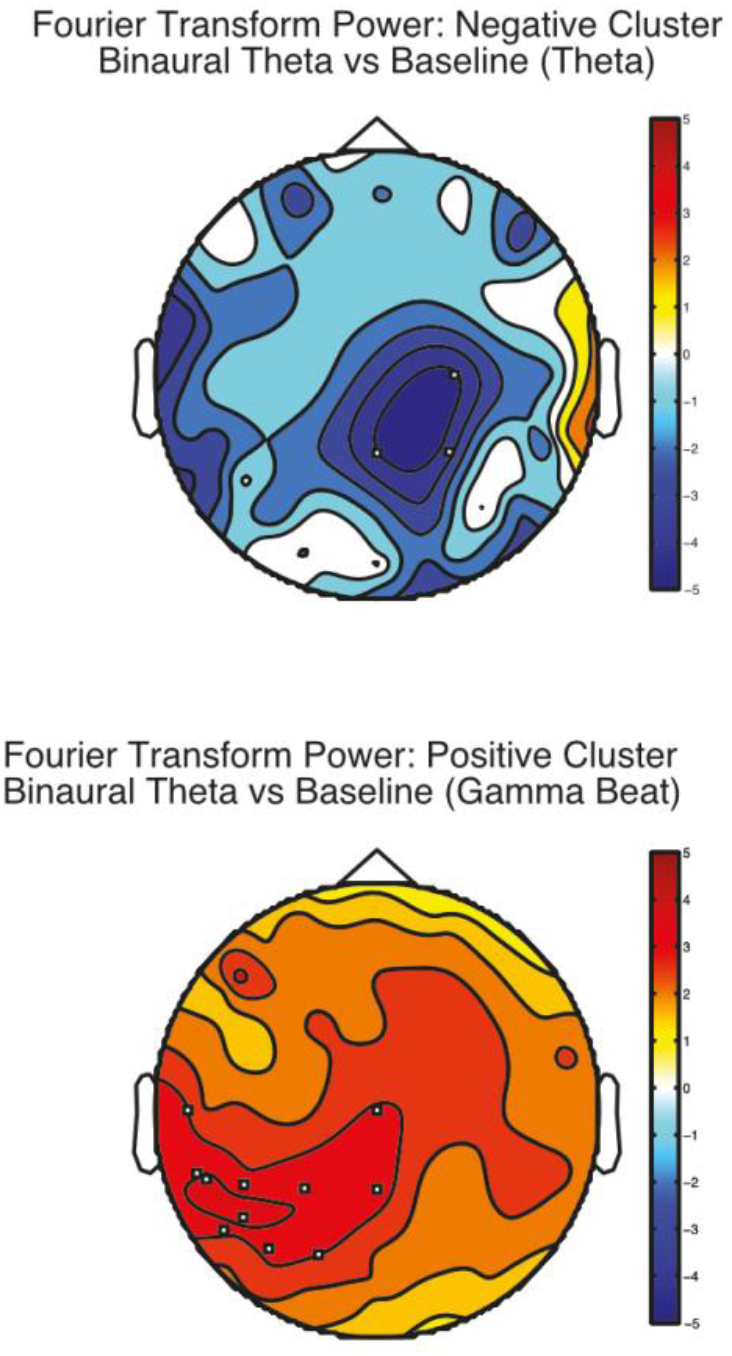
Fourier transform power used as a local synchronization index in Theta conditions. Topographies were averaged across participants and compared to baseline. Fourier Transform Power was normalized using z scores. We only show contrasts that exhibit at least three significant electrodes here (depicted as small white squares). Non-significant contrasts can be found in the supplemental material (refer to **Figure S4** (Theta conditions) and **Figure S5** (Gamma conditions)). Frequency band limits are as follows: Delta (1 - 4 Hz), Theta (5 - 8 Hz), Alpha (9 - 12 Hz), Beta (13 - 30 Hz), Gamma (32 - 48 Hz), Theta Beat (6 - 8 Hz) and Gamma Beat (39 - 41 Hz). The color bar indicates T-values from student’s test. Please note that no iCOH contrasts were significant.

#### Neurophenomenological analysis

When taking into consideration individual differences in terms of subjective experience, we find neural connectivity patterns associated with high Absorption Depth and Mental Relaxation that are consistent across participants. When contrasting each participants’ two highest and two lowest rated experimental conditions in terms of mental relaxation, we found one negative frontal cluster of local activity (**Hilbert Transform, Figure 6a**) at Theta frequency band (CS = −9.05, p = 0.028), and two negative long-range clusters of activity (**both PLV and iCOH**) at the same frequency band: a frontal to occipital connectivity pattern (**Figure 6c**, PLV, CS = −77.49, p = 0.058) and a frontocentral to occipital one (**Figure 6d left** iCOH: CS = −61.95, p = 0.041). Finally, we found a frontal to right-temporal-occipital negative cluster of activity at 40 Hz (**Figure 6d right** iCOH: CS = −97.325326, p = 0.045). On the other hand, we contrasted the two top experimental conditions in which a given participant rated absorption depth the highest with the experimental conditions in which they rated absorption the lowest. We find three negative clusters of local (**Fourier Power**) activity at 40 Hz (**Figure 6b**): a left temporal-occipital one (CS = −9.77, p = 0.041000), a central one (CS = −10.03, p = 0.04) and a right temporal one (CS = −10.168336, p = 0.035000). All neurophenomenological contrasts (both significant and non-significant) can be found in the supplemental material (**Figure S6** for PLV and Hilbert Amplitude, and **Figure S7** for iCOH and Fourier Power).

**Figure 6.**
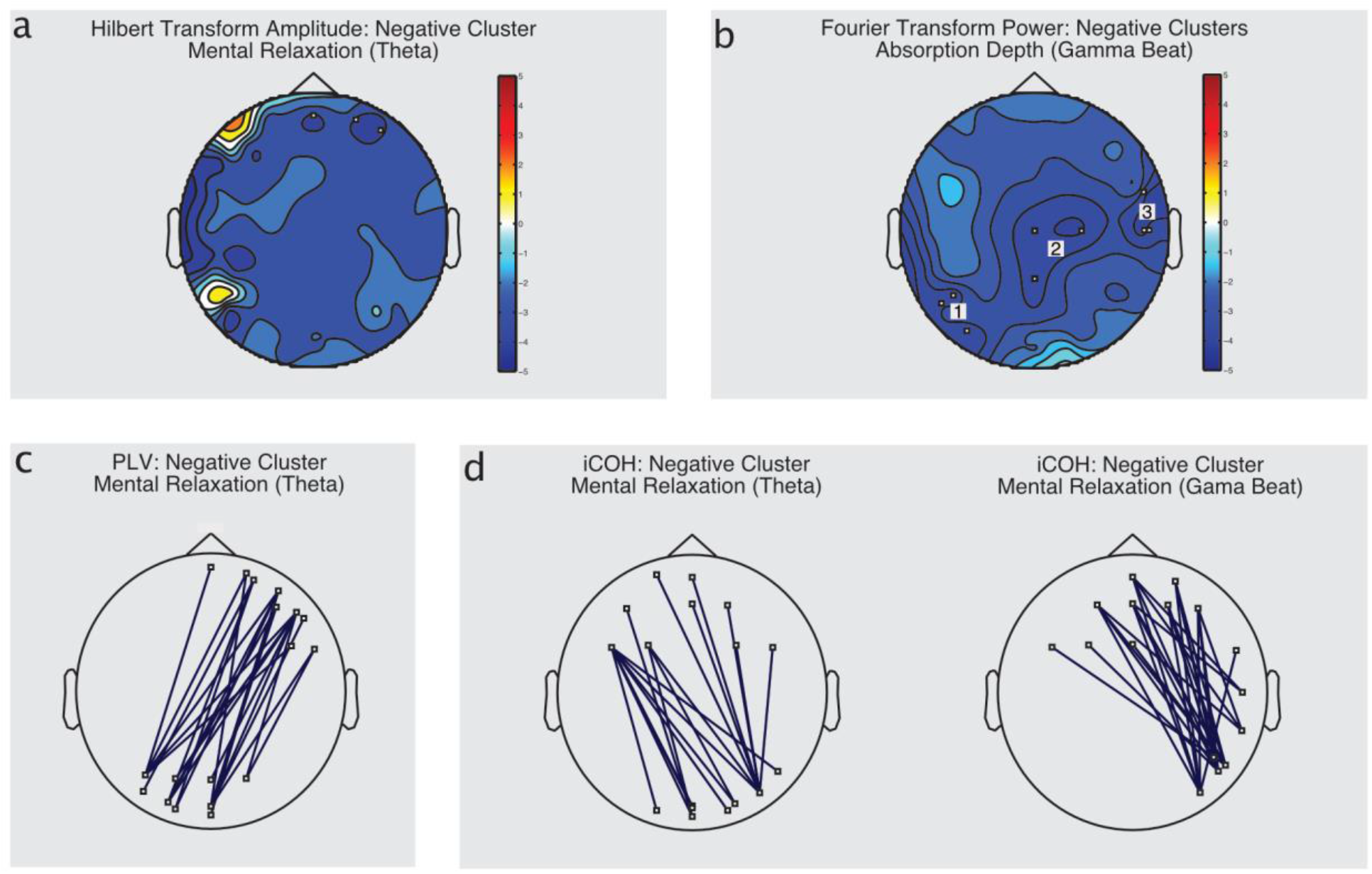
Neurophenomenological analysis: correlates between subjective experience and EEG connectivity patterns. Each participants’ two highest rated (Mental Relaxation and Absorption Depth) experimental conditions (binaural gamma, monaural theta…) were contrasted with the two lowest rated conditions. These contrasts were then averaged across participants for each separate scale (Mental Relaxation and Absorption Depth). All statistics were normalized using z scores. We only show contrasts that exhibit at least three significant electrodes here (depicted as small white squares). All contrasts can be found in the supplemental material (refer to **Figure S6** (PLV and Hilbert Transform Amplitude) and **Figure S7** (iCOH and Fourier Transform Power)). Frequency band limits are as follows: Delta (1 - 4 Hz), Theta (5 - 8 Hz), Alpha (9 - 12 Hz), Beta (13 - 30 Hz), Gamma (32 - 48 Hz), Theta Beat (6 - 8 Hz) and Gamma Beat (39 - 41 Hz). **(a) Hilbert transform amplitude used as a local synchronization index.** The color bar indicates T-values from student’s test. **(b) Fourier transform power used as a local synchronization index.** The color bar indicates T-values from student’s test. **(c) Phase locking value used as an index of long-distance synchronization between electrodes.** Blue lines indicate a significant negative phase locking value (PLV) between two electrodes. **(d) Phase locking value used as an index of long-distance synchronization between electrodes.** Blue lines indicate a significant negative phase locking value (iCOH) between two electrodes.

## Discussion

Here, we asked whether binaural beats are able to elicit neural entrainment, and modulate mood, in a specific fashion compared to a sham rhythmic stimulus. To do so, we used a passive, single-blind listening task where participants were exposed to both binaural and sham conditions while we recorded their electrical brain activity and self-reports related to mood. By comparing activity between binaural and sham conditions at different levels (subcortical, cortical and self-reports), we found that binaural beats entrained the cortex, but the sham condition did so more strongly, with none of them showing an effect on mood. Furthermore, while distinct Functional Connectivity patterns emerged for both binaural and monaural beats at different frequencies, these are not consistent with previous literature and are not related to participants’ self-reported mood.

### Binaural and monaural beats elicit subcortical responses at carrier frequencies

Though it is commonly agreed that Binaural Beats originate in the brainstem (Oster, 1973; Moore, 2012; Wernick & Starr, 1968), to the best of our knowledge, we are the first ones to investigate this particular stimulus at subcortical levels using EEG. As we predicted, both experimental conditions (theta and gamma), regardless of beat type, elicited a frequency following response at the pure tone frequencies, with no difference between monaural and binaural beats. This is consistent with the existing auditory brainstem response literature, where the generated subcortical responses are found to have a close spectrotemporal structure to the patterns of an acoustic stimulus, such as speech syllables (Skoe & Kraus, 2010; Lehmann & Schönwiesner, 2014). Furthermore, given our choice of carrier frequencies (around 400 Hz), it is very unlikely these responses have a cortical origin (see Coffey et al., 2016). The lack of difference between beat types suggests that both stimuli are being processed in the same way at the subcortical level.

### Monaural beats elicit higher cortical entrainment at the beat frequency than their binaural counterparts

Both beat types entrained the brain at their beat frequencies, with monaural conditions eliciting the highest response when compared to binaural conditions. In terms of Theta beat frequency, both Jirakittayakorn & Wongsawat (2017) and Karino et al. (2006), found similar entrainment using Theta binaural beats with exposure times between five and ten minutes. Following Garcia-Argibay’s conclusions (2018), relatively long exposure time and non-masked binaural beats (i.e. not using white or pink noise to mask them) seem to optimize the responses to the beats. In terms of Gamma beat frequency, we successfully replicated previous studies (Schwarz & Taylor, 2005; Draganova et al., 2007; Ross et al., 2014): both binaural and monaural gamma beats entrain cortical activity at 40Hz, but binaural beats elicit less power at the beat frequency. One possible explanation as to why binaural beats elicit less power than monaural beats is that the entrainment we measure at the cortical level might be caused by the perceived rhythmicity, and not the binaural beat itself. Subjects’ tended to report that the modulation (i.e. the beat) intensity in binaural beats was weaker than that of the monaural beat. The ASSR correlates with stimulus’ loudness (Van Eeckhoutte et al., 2016; Picton et al., 2007; Lins & Picton, 1995), which might explain the difference in ASSR power in the frequency domain. Furthermore, we root mean squared (rms) normalized our stimuli, precluding loudness as an explanatory factor for the difference in ASSR power. Using proper statistical and experimental control, and contrary to popularized marketing claims, here we show that binaural beats weakly entrain the cortex when compared to other rhythmic stimuli, such as monaural beats.

### Binaural and monaural beats fail to modulate mood

Echoing previous reports, we did not find evidence of binaural beats, nor monaural beats, modulating cognitive states or mood: López-Caballero & Escera (2017) found no emotional regulation due to binaural beats as indexed by changes in heart rate and skin conductance, while Gálvez et al. (2018) found no modulation of anxiety as indexed by the State Anxiety Inventory (SAI). This stands in contrast with other reports where cognitive performance and mood were successfully modulated by binaural beats (Wahbeh et al., 2007, Le Scouarnec et al., 2001, Padmanabhan et al., 2005; Isik et al., 2017, Reedijk et al., 2013; Reedijk et al., 2015, Garcia-Argibay et al., 2018).

### Both beats elicit differential short-range connectivity patterns

Monaural and binaural beats affect short range electrode level connectivity patterns differentially. We only find a significant short-range effect in monaural gamma and binaural theta, suggesting both beat type and beat frequency are relevant and precluding conclusions such as this activity being a by-product of sustained listening or binaural integration. Our gamma findings are in accordance with those from Becher et al. (2015). Using both intracranial and scalp EEG, they found peak EEG power at 40 Hz (gamma beat) at the scalp electrodes using the power of the envelope of the signal. They found a similar effect in temporo-lateral intracranial electrodes, suggesting this entrainment originates in auditory cortices. Furthermore, they found a significant decrease in EEG power at 5 Hz (theta frequency) in temporo-basal anterior and posterior areas, which might explain the local activity in our participants. This activity could be in line with a dipole from auditory cortices pointing upwards, which suggests there is only one active cortical source. On the other hand, binaural theta conditions elicited a positive parietal cluster at 40 Hz (see cross frequency section). The functional meaning (if any) of this short-range patterns remains unclear as we found no difference in participants’ self-reports.

### Both beats elicit long-range connectivity patterns indexed only by PLV

To investigate long-range connectivity, we used two different but complementary statistics: the imaginary part of coherence and the phase locking value. iCOH gets rid of all interactions that have zero to very small time delays, while the PLV quantifies how consistent phase differences are between electrodes. We only found Functional Connectivity patterns indexed by PLV. Because of this, it is unclear whether these patterns are due to one source being propagated around the scalp, or if there are multiple sources active with an almost zero-time delay between them (Nolte et al., 2004). We see differential effects between beat types and frequencies, as well as cross-frequency interactions (discussed in the next section). Gamma experimental conditions where the only ones to elicit within frequency activity: Monaural gamma elicited a cluster of scalp-wise connectivity that is not consistent with previous research. Using intracranial electrodes, Becher et al. (2015) showed phase desynchronization at mediotemporal areas using a 40 Hz monaural beat, whereas we found scalp-wise synchronization using a very similar stimulus. Furthermore, Schwarz et al. (2005) showed that there was a delay of several milliseconds in the activity elicited across the fronto-occipital axis, using a 40 Hz monaural beat, suggesting multiple cortical sources of activity. Because the iCOH analysis did not reveal significant connections between electrodes, it is unclear whether the phase differences Schwarz et al. (2005) report are due to volume conduction or the connectivity patterns we found are caused by multiple, but tightly synchronized, brain regions. Because we did not find any difference in subjective reports, the functional meaning (if any) of this activity remains unclear.

### Binaural beats elicit cross-frequency connectivity patterns

Binaural theta conditions elicit a front to back, cross-frequency connectivity pattern at gamma beat frequency (40 Hz) while binaural gamma elicits a widespread connectivity pattern at alpha frequency. In line with our results, several groups have found binaural beats eliciting activity outside of the frequency range of the beat. Despite this, these findings do not seem to be consistent: using theta binaural beats, Gao et al. (2014) found a decrease in relative beta power over left temporal areas and Ioannou et al. (2015) report no significant difference from theta binaural beats at other frequency bands.

We found that binaural theta beats elicit activity at higher frequencies (40 Hz), while binaural gamma beats elicit activity at lower frequencies. This cross-frequency coupling (low frequency driving a higher frequency and a higher frequency driving a lower frequency) could be evidence for large-scale integration being enhanced by binaural beats. Varela et al. (2001) argue that slower rhythms (such as theta) provide a temporal framing for faster oscillations. For example, gamma oscillations are thought to leverage this slower temporal framing during successive cognitive moments of synchronous assemblies where memory is consolidated (Osipova et al., 2006; Burke et al., 2013; Lisman & Jensen, 2013). We did not investigate cognitive processes directly, but there is evidence that binaural beats impact memory as a function of beat frequency—beta frequencies seems to enhance it (Lane et al., 1998; Kennerly et al., 1994; Beauchene et al., 2016; Gálvez et al., 2018) while theta frequencies have an inconsistent effect (negative in some cases: Garcia-Argibay et al., 2017, Beauchene et al., 2016; and positive in others: Ortiz et al., 2008). In our specific experiment, our stimuli failed to modulate mood as self-reported by participants, but these cross-frequency interactions might provide a framework explaining why binaural beats are able to modulate cognitive performance in other reports.

### Individual differences shed light on connectivity patterns associated with specific cognitive states

We found a consistent pattern of deactivation and desynchronization related to participants’ self-reports of Mental Relaxation and Absorption Depth. High Mental Relaxation is associated with theta frequencies in a frontal cluster of local desynchronized activity, with a front to back desynchronization of activity at both theta and gamma indexed by both PLV and iCOH. This converging evidence suggests this activity is robust and not due to volume conduction. Absorption Depth, on the other hand, is associated with three clusters of activity: one in the center of the scalp, and one on each side of the head (around temporal and parietal areas). Changes in the anterior and frontal midline in theta power have been related to emotionally positive states (Aftanas & Golocheikine, 2001), and meditation-related states (Baijal & Srinivasan, 2010). Baijal & Srinivasan (2010) found a similar deactivation pattern in parietal and occipital areas accompanied by frontal theta activation associated with meditative states. In line with Travis & Wallace’s (1999) model of Transcendental Meditation, our results could be interpreted as frontal, central, and parietal activity driven by basal ganglia and thalamocortical structures that regulate “quieter levels of cortical functioning”. On the other hand, Hinterberger et al. (2014) found similar central and parietal gamma deactivation patterns during meditative tasks. Taking all this information together, our Functional Connectivity results point at a state similar to meditation characterized by heightened Mental Relaxation and Absorption Depth. Despite this, we were not able to relate this specifically to any of our experimental conditions.

### Limitations and Future Directions

Though several of our findings seem to be consistent with existing literature, we acknowledge that we only recruited 16 people and that these findings should be replicated with higher sample sizes. We instructed participants to close their eyes during the whole experiment, which might have not been ideal, especially because a couple of participants reported high drowsiness and two reported falling asleep. Future binaural beats studies should look at different ways of indexing connectivity at both scalp and source level (Solcà et al, 2016). The study of binaural beats will also greatly benefit from the transition of a mass univariate statistical framework (such as the one used here) to a multivariate statistical framework (see McIntosh & Mišić, 2013). Fields such as Graph Theory present promising opportunities to determine the characteristics of cortical networks and summarizing large numbers of data points into a few statistics to truly understand how binaural beats affect the brain (Ioannou et al., 2015; Ala et al., 2018). On a more technical note, we did not alternate the polarity of the stimulus of the binaural beats (a common practice in the Auditory Brainstem Response literature), which might have affected the brainstem responses we found. Nevertheless, the transducer with which the stimuli were presented to the participants was magnetically shielded, a procedure that is known to minimize stimulus artifacts (Skoe & Kraus, 2010).

The study of binaural beats has been around for a number of years. Despite this, few advances have been made in the field. As Garcia-Argibay et al. (2018) concluded, there are several mediating variables—such as beat frequency, exposure time or stimulus masking—that are not always clearly reported. Furthermore, we conclude that there are two important factors that are usually considered as trivial: proper EEG analysis and the use of “sham” conditions. Future binaural beats studies should be mindful of these variables and report them accordingly. Several studies that did not find any entrainment to the beat frequency of binaural beats (e.g., López-Caballero & Escera, 2017) did not use standard normalization practices (for an in-depth discussion of these procedures, see Cohen, 2014, Ch. 18). The human EEG spectrum exhibits a 1/f power scaling (similar to pink noise). By properly normalizing data using a baseline condition, we ensure that all data (i.e. the different frequency bands) will have the same scale, and we are appropriately disentangling background and task-unrelated dynamics (Cohen, 2014). Researchers should be mindful of the analysis approach they are taking, as well as using an appropriate “sham” condition to truly elucidate whether binaural beats are a special kind of stimulus or their advantages are due to stimulus properties (such as the rhythmicity in the signal).

Future studies should carefully choose exposure time, the performance or mood measurement and the frequency of the beat. As Garcia-Argibay et al. (2018) concluded, higher exposure times are associated with larger effect sizes. Nevertheless, whether several sessions will present an increased entrainment and performance/mood boost, and whether there are carryover effects that are sustained even after stimulation ceases, are still open questions. Binaural beats seem to modulate memory and attention performance, as well as anxiety and analgesia. Our findings provide a plausible base explanation as to why memory and attention performance could be modulated by binaural beats (i.e. binaural beats elicit cross-frequency interactions). Future studies should focus on measuring cognitive performance on both attention and memory tasks, and mood regulation related to anxiety and pain perception. Participants should be exposed to the stimulation both before and after the task (Garcia-Argibay et al., 2018). Finding relations between the degree of cross-frequency modulations and cognitive performance in attention and memory tasks will help us better elucidate whether binaural beats can be used for cognitive enhancement or not. Finally, the choice of frequency is not trivial: alpha, beta, and gamma seem to provide positive effects in memory tasks, while theta frequency seems to hinder it in most cases (Garcia-Argibay et al., 2018).

## Conclusions

Using a factorial experimental design and a single-blind, passive listening task, we aimed to elucidate the impact of binaural beats on the brain. We found that a sham condition was better at synchronizing brain activity than binaural beats and, though both elicited differential patterns of connectivity, they did not modulate mood. Contrary to marketing claims, we failed to find evidence for binaural beats being a “special kind of stimuli” that modulates mood and entrains the brain in a specific fashion. Future studies should look at cognitive performance modulation (especially with attention and memory tasks) using binaural beats with multi-session approaches to elucidate if they elicit permanent effects. This line of research has important implications: marketing claims from companies commercializing binaural beats may be based purely on placebo effects. By using a neuroscientific lens with statistical and scientific rigor at its core, we can study “alternative” mood regulation practices to ensure the general public makes informed decisions. All in all, binaural beats weakly entrain the cortex and elicit connectivity patterns that were not elicited by a “sham” stimulation. Whether these connectivity patterns have a functional meaning (in terms of cognitive enhancement and mood modulation) or not, is still an open question.

## Acknowledgements

The authors of this paper would like to thank Pierre Rainville and Bérangère Houze for sharing the E-SAS scales; Mihaela Felezeu for all the help and support during data acquisition; and our participants for volunteering their time to perform the experiment.

