## Supplemental figures for "Binaural Beats through the auditory pathway: from brainstem to connectivity patterns"

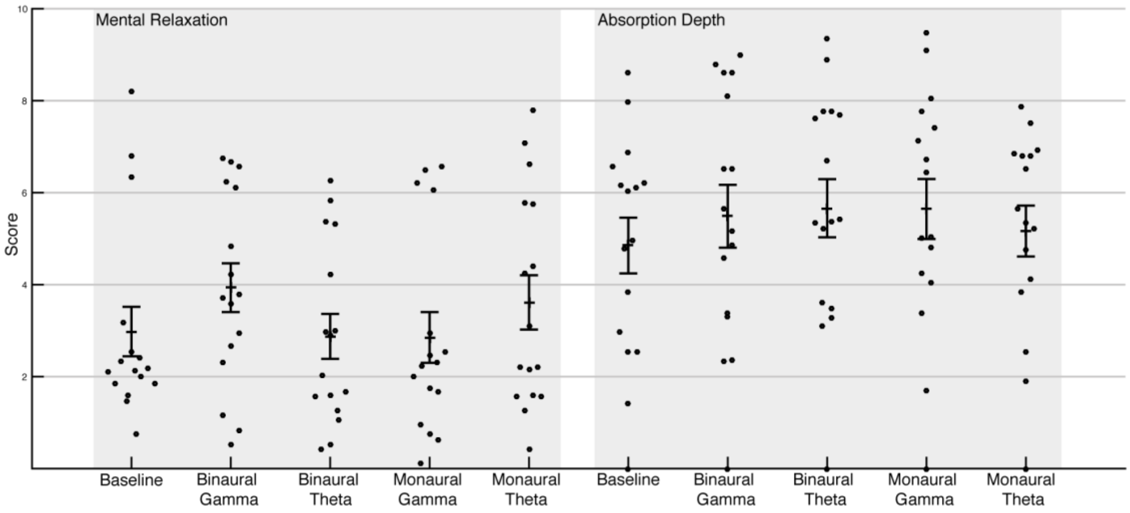


**S1. Visual Analogue Scales: Mental relaxation and absorption depth.** Each data point represents one participant's score. Mean and standard error of the mean are plotted on top of individual data points.

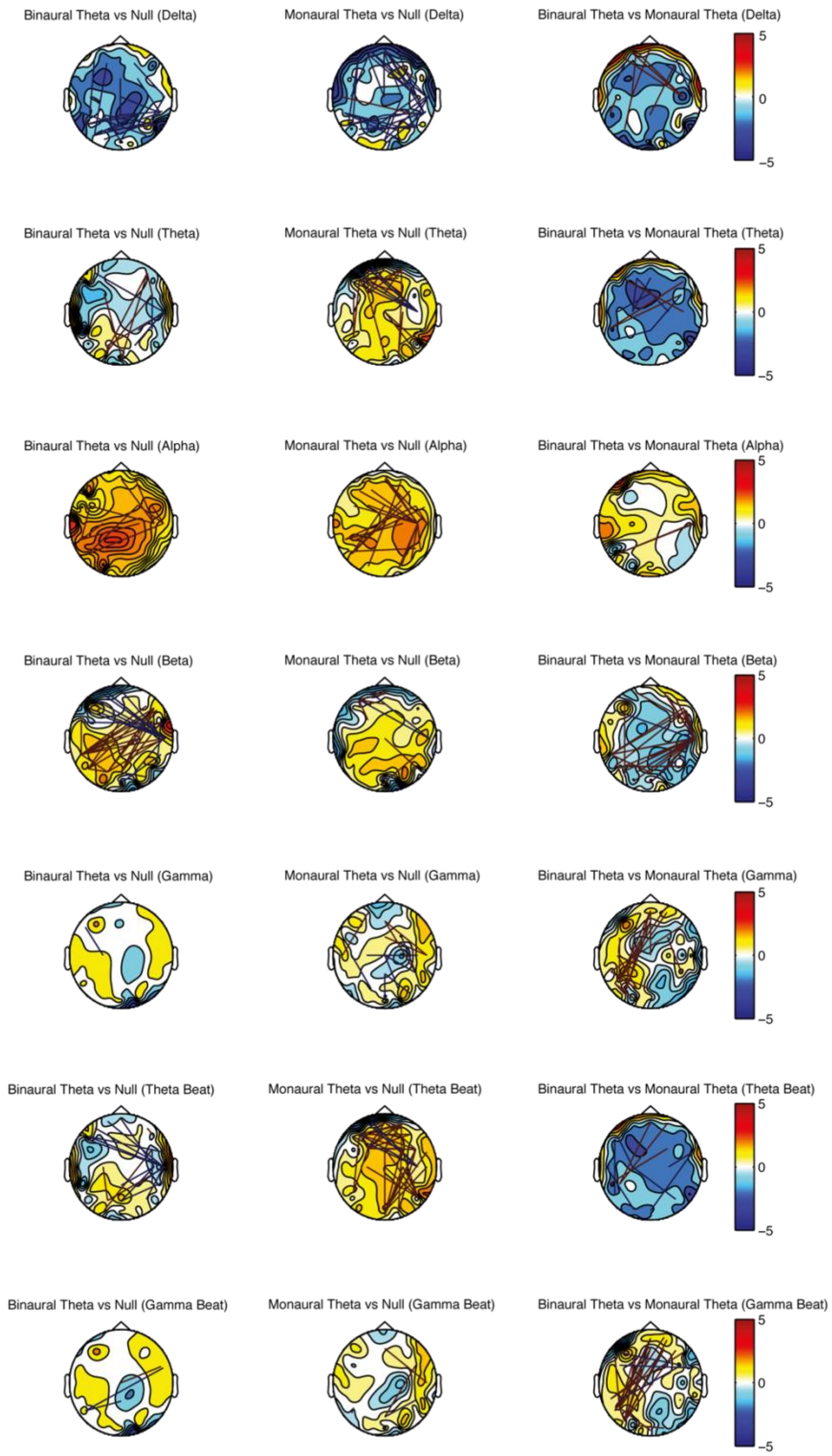


**Figure S2. Contrast topographies for Phase-locking value (PLV) and Hilbert Transform amplitude during theta beat conditions.** See main text for details.

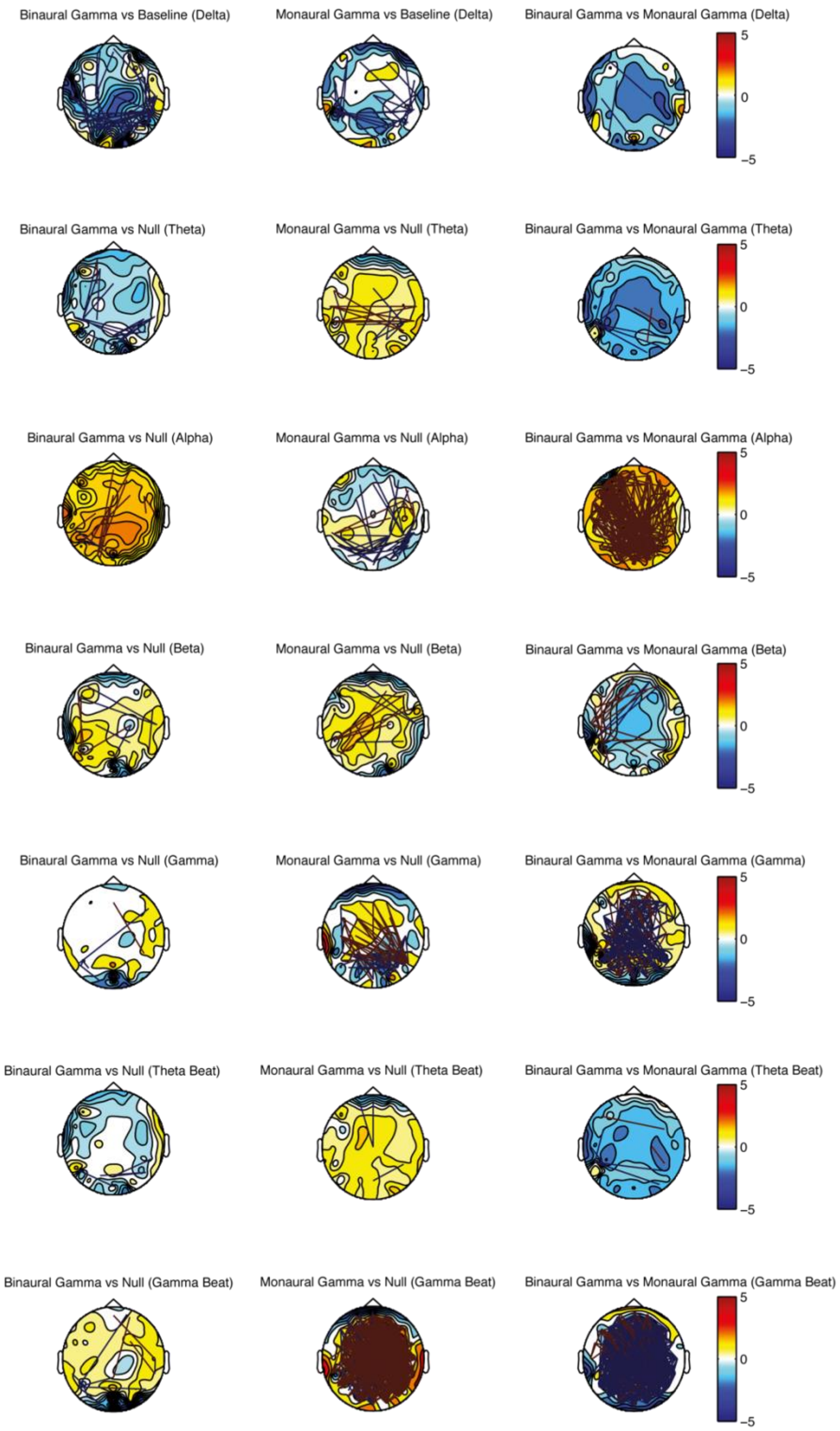


**Figure S3. Contrast topographies for Phase-locking value (PLV) and Hilbert Transform amplitude during gamma beat conditions.** See main text for details.

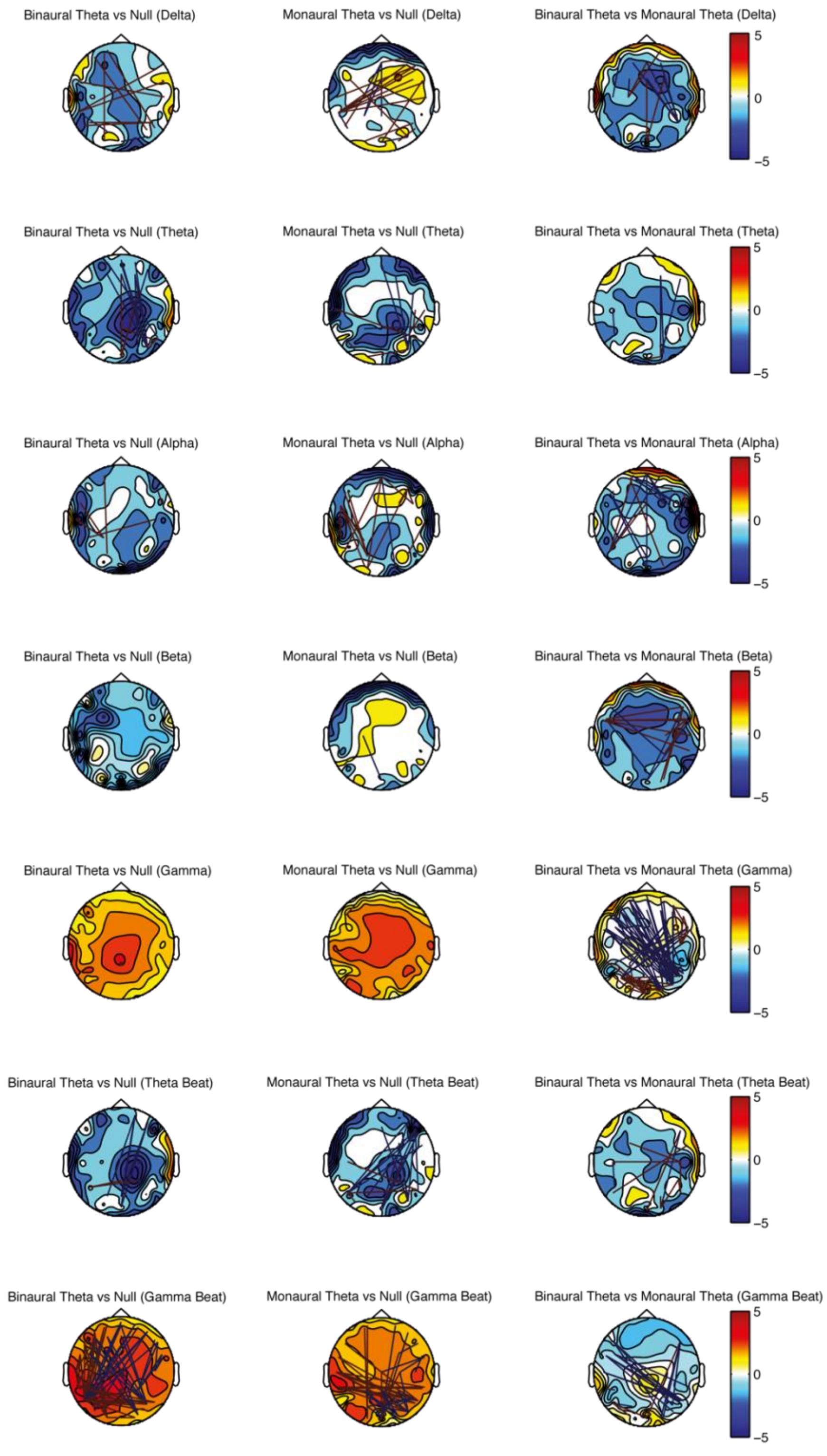


**Figure S4. Contrast topographies for imaginary coherence (iCOH) and Fourier Transform power during theta beat conditions.** See main text for details.

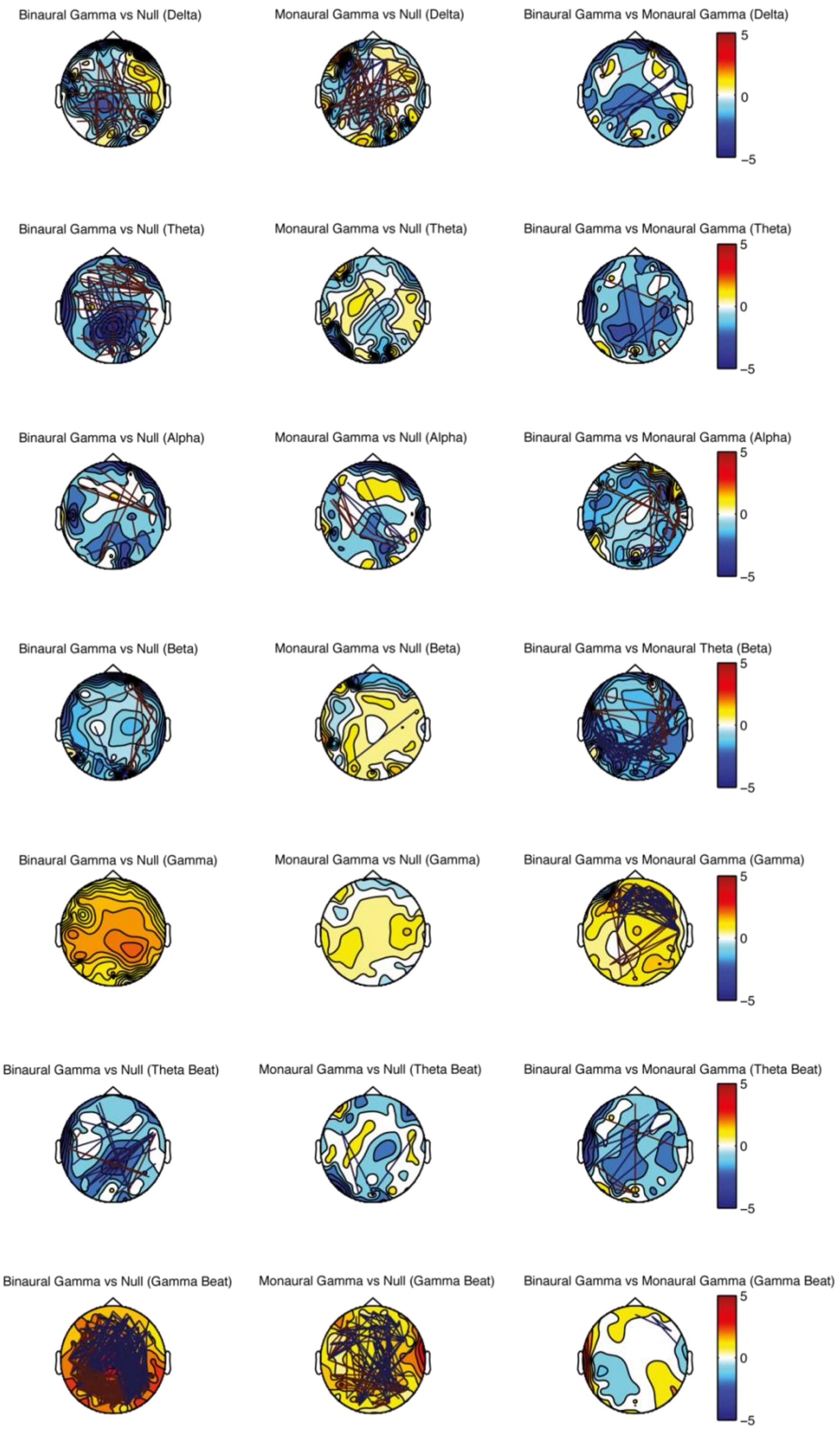


**Figure S5. Contrast topographies for imaginary coherence (iCOH) and Fourier Transform power during gamma beat conditions.** See main text for details.

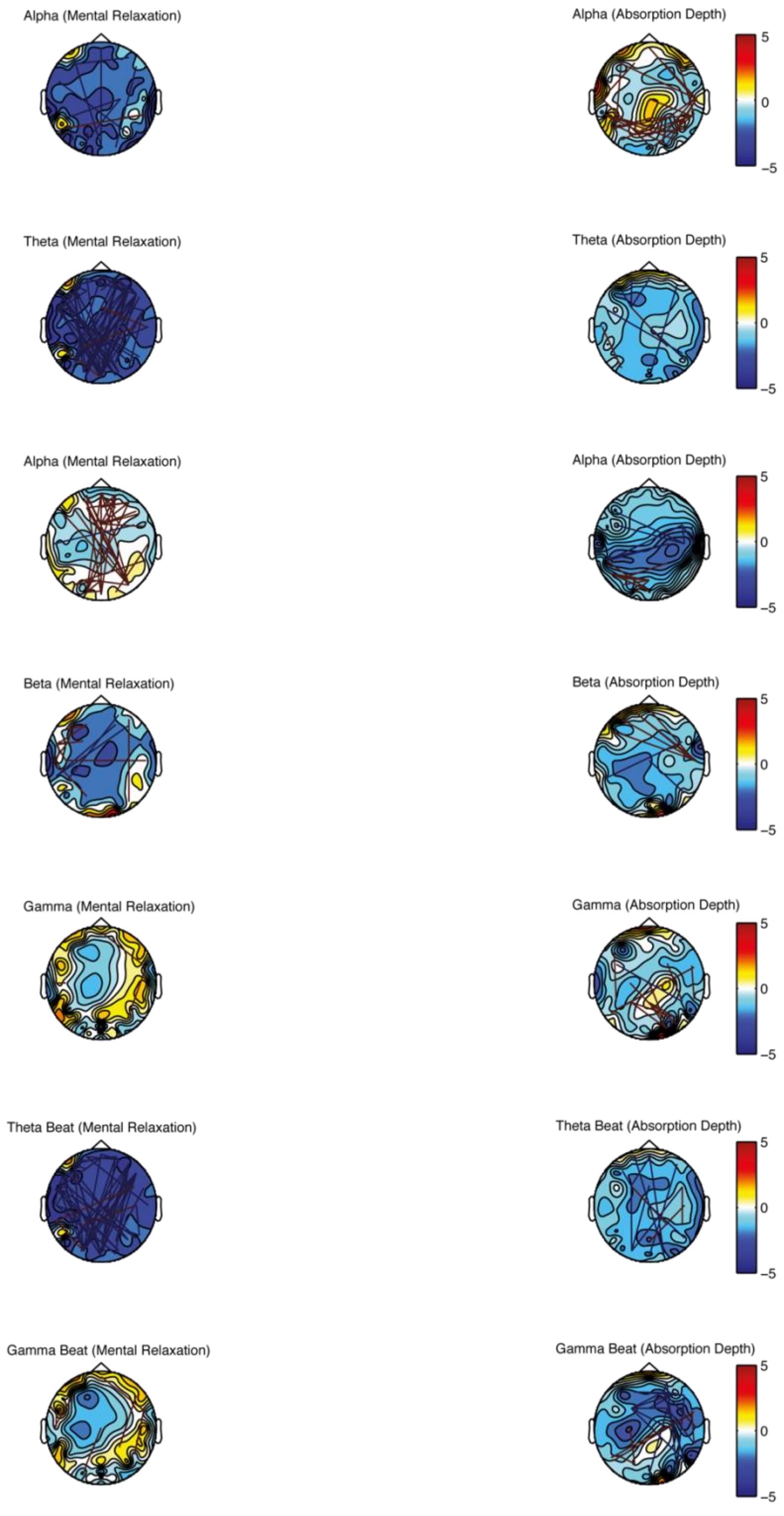


**Figure S6. Neurophenomenological analysis: Contrast topographies for Phase-locking value (PLV) and Hilbert Transform amplitude.** See main text for details.

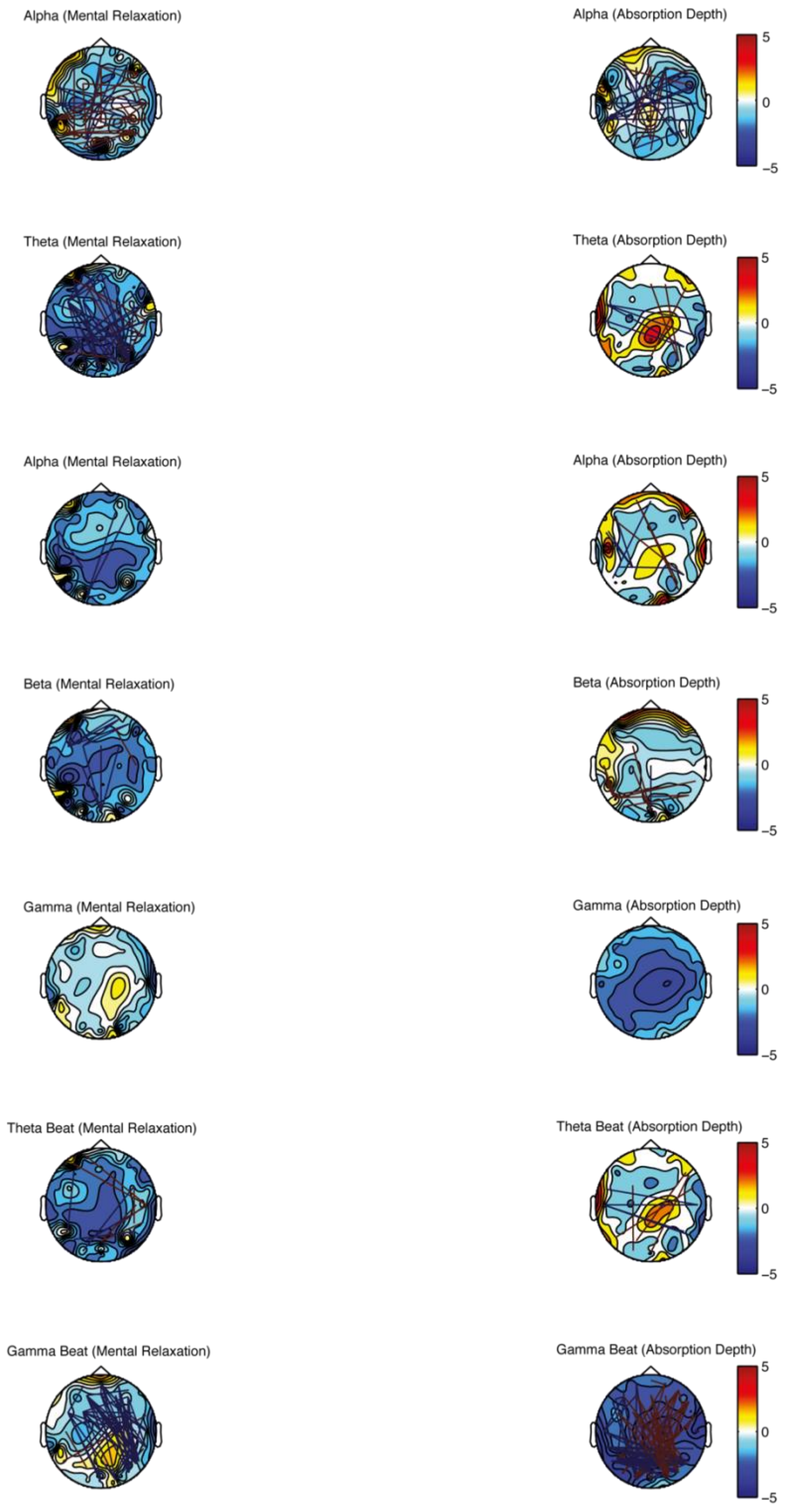


**Figure S7. Neurophenomenological analysis: Contrast topographies for imaginary coherence (iCOH) and Fourier Transform power.** See main text for details
